# Common synaptic inputs and persistent inward currents of vastus lateralis motor units are reduced in older age

**DOI:** 10.1101/2023.02.03.526933

**Authors:** Yuxiao Guo, Eleanor J. Jones, Jakob Škarabot, Thomas B. Inns, Bethan E. Phillips, Philip J. Atherton, Mathew Piasecki

**Affiliations:** Centre of Metabolism, Ageing & Physiology (COMAP), MRC-Versus Arthritis Centre for Musculoskeletal Ageing Research & National Institute for Health Research (NIHR) Nottingham Biomedical Research Centre, School of Medicine, University of Nottingham, Derby, UK; School of Sport, Exercise and Health Sciences, Loughborough University, Loughborough, UK

**Keywords:** Ageing, motor unit, persistent inward current, common synaptic input, vastus lateralis

## Abstract

Although muscle atrophy may partially account for age-related strength decline, it is further influenced by alterations of neural input to muscle. Persistent inward currents and the level of common synaptic inputs to motoneurons influence neuromuscular function. However, these have not yet been described in aged human quadriceps.

High density surface electromyography (HDsEMG) signals were collected from the vastus lateralis of 15 young (mean±SD, 23 ± 5 y) and 15 older (67 ± 9 y) men during submaximal sustained and 20-s ramped contractions. HDsEMG signals were decomposed to identify individual motor unit discharges, from which delta F and intramuscular coherence were estimated.

Older participants produced significantly lower knee extensor torque (p<0.001) and poorer force tracking ability (p<0.001) than young. Older participants also had lower delta F (p=0.001) and coherence estimates in the alpha frequency band (p<0.001) during ramp contractions when compared to young.

Persistent inward currents and common synaptic inputs are lower in the vastus lateralis of older males when compared to young. These data highlight altered neural input to the clinically and functionally important quadriceps, further underpinning age-related loss of function which may occur independently of the loss of muscle mass.

**Key Points:**

- The age-related loss of muscle mass is exceeded by the loss of function, which is influenced by structural and functional alterations of the nervous system.
- Motoneuronal persistent inward currents and common synaptic inputs play an important role in the activation of motor units and subsequent force generation and control ability.
- Here we show reduced estimates of persistent inward currents and lower common synaptic inputs to older vastus lateralis, potentially contributing to observed lower strength and poorer force tracking.
- These findings highlight decrements of the aged human motor system, accompanied by muscle atrophy in functionally relevant muscle groups, which should be considered in the application of interventions targeting aged human muscle.

## Introduction

Human ageing is characterised by a progressive reduction in muscle size and muscle strength, resulting in an impairment of force generation, force control and physical performance [1–3], all of which can negatively impact quality of life [4,5]. The loss of muscle size is explicable by the combined effects of muscle fibre atrophy and muscle fibre loss [2], and although atrophy may partially account for declines in muscle strength, it is further influenced by structural and functional alterations of the nervous system.

The voluntary contraction of muscle fibres occurs via net excitatory signals to the motor units (MUs); groups of muscle fibres innervated by a single motoneuron [6]. The central nervous system controls muscle force via two primary strategies; varying the number of MUs recruited and varying the discharge rate of each motor neuron, both of which are susceptible to age-related alterations. Substantial motoneuron loss accelerates in the sixth decade of life, resulting in fewer MUs, and a compensatory process of MU remodelling further results in larger MU innervation ratios [7,8]. Additionally, although this appears to be muscle specific, an age-related decrease in MU discharge rate has also been documented at normalised contraction levels [9,10] as well as being associated with circulating hormone levels that are also affected by age [11].

In addition to direct neurotransmitter activation onto postsynaptic ligand-gated channels at the spinal cord level via ionotropic mechanisms, activation of the motor nervous system also relies on neuromodulation [12]. This is mediated by intracellular second messenger systems that modify the voltage- and ligand-gated channels of neurons and alters several properties from their level of excitability to their pattern of firing in response to a given input [13,14]. This process is enhanced by monoamines, e.g. serotonin and norepinephrine released from the brainstem nuclei by stimulating inward-flowing persistent calcium and sodium currents [12], known as persistent inward currents (PICs). As long as the membrane potential remains above the activation threshold, these PICs tend to remain activated [12], combined with neuromodulatory drive amplifying the firing behaviours of motoneurons to synaptic inputs up to five fold [15] as well as generating long-lasting plateau potentials which results in self-sustaining firing [16]. Even though PICs appear to decrease with age in upper [17] and lower limb muscles [18], PICs can be mediated by local ionotropic inhibitory inputs that may be muscle specific in the response to age [10,19,20], particularly those highly susceptible to functional losses such as the vastus lateralis (VL) [21].

Rather than controlling the discharge pattern of individual MUs, the central nervous system controls the excitatory inputs to the MU pool, referred to as common synaptic inputs [22] which are defined as the proportion of net synaptic input correlated between MUs. Force fluctuations are largely determined by the alterations in low frequency components of common synaptic inputs to MUs, which is reflected in concurrent fluctuations in MU discharge rates from the same MU pool [23]. Accordingly, a decrease in force steadiness associated with ageing is a consequence of an increase in variance of common synaptic inputs [24,25]. However, the variability of common synaptic inputs in quadriceps has only been explored in healthy young individuals [26,27], with no available data from older humans.

It is not possible to make intracellular recordings from human spinal motoneurons in order to estimate PICs, so alternative methods are employed which distinguish intrinsic excitability of the MU from its descending synaptic drive by using the unit-wise MU analysis technique [17]. In view of the advances with which MU discharging patterns can be measured, it provides an opportunity to investigate and better understand the physical properties and the underlying mechanisms which may impair motor control in older age. Therefore, the aim of this study was to explore the differences in physical performance as well as the magnitude of persistent inward currents and common synaptic inputs in the vastus lateralis between healthy young and older males. We hypothesized that older participants would exhibit greater physical decrements and lower levels of persistent inward currents and common synaptic inputs.

## Methods

### Participants and ethical approval

This study was approved by the University of Nottingham Faculty of Medicine and Healthy Science Research Ethics Committee (90-0820, 199-0221) and conformed to the Declaration of Helsinki.

15 healthy young males between the ages of 18–40 (mean ± SD; age: 23.1 ± 5.2 years; body mass index: 25.0 ± 2.5 kg/m^2^) and 15 healthy older males between the ages of 55 to 85 (67.2 ± 8.9 years; 26.1± 2.3 kg/m^2^) provided informed consent to take part in the study. All participants were recruited through advertisements in the local community. Prior to enrolment, all participants completed a comprehensive clinical examination and metabolic screening was conducted at the School of Medicine, Royal Derby Hospital Centre. All recruited participants were recreationally active and would be excluded if they display evidence of: BMI < 18.5 or > 35kg/m^2^; are competitive in an athletic discipline at county level or above; musculoskeletal disorders; respiratory disease; neurological disorders; metabolic disease; active cardiovascular problems; active inflammatory bowel or renal disease; recent steroid treatment within 6 months or hormone replacement therapy; family history of early (< 55years) death from cardiovascular disease.

### Anthropometry

Ultrasound was used to measure the cross-sectional area (CSA) of the VL muscle of the right leg, using an ultrasound probe (LA523 probe, B-mode, frequency range 26– 32Hz, and MyLabTM50 scanner, Esaote, Genoa, Italy) at the anatomical mid-point of the right thigh. Images were recorded with the participants laying in a supine position and the mid-point obtained from the identification of the greater trochanter at the top of the femur and the mid-point of the patella. Panoramic imaging (VPAN) images were obtained in a medial to lateral fashion, beginning and ending the capture of the image at the aponeurosis borders of the VL of the right leg. The produced images were subsequently analysed using ImageJ software (National Institute of Health, USA) through tracing around the contour of the aponeurosis via polygon function to quantify the area of the VL. Each image was analysed three times and the mean value of three images was accepted as CSA.

### Experimental Protocol

All participants were required to attend the lab at 0900 after an overnight fast and abstain from strenuous exercise 24h prior to the testing sessions. Participants were seated in a custom-built chair with hips and knees flexed at ∽90 degree. The lower leg was secured to a force dynamometer with noncompliant straps above the medial malleolus. To minimize the movement of the upper trunk during testing, a seat belt was fastened across the pelvis of the participant. After a standardised warm up through several submaximal contractions [28], participants were instructed to contract as hard and fast as they could. During the trial, participants were not allowed to hold onto the side of the chair and were asked to cross their arms across the chest. This was performed with real-time visual force feedback on a monitor placed in front of the participant, with verbal encouragement to aid in producing maximal effort. This was repeated two to three times further, giving 60 seconds of rest between attempts standardised recovery. If the difference between the last two attempts was < 5%, the highest value recorded in newtons was taken and accepted as the maximal isometric voluntary contraction force (MVC). Muscle torque was calculated by multiplying MVC by the lever arm length from the knee to the centre of the ankle strap.

Following a 60-second rest, participants were then asked to perform four submaximal voluntary isometric contractions at 25% of MVC (∽12s) and four triangular shaped contractions (10s up and 10s down) peaking at 20% of MVC, with 30-second intervals between contractions. In each case, a target line was displayed on a screen and participants were instructed to follow the target as close as possible. Ramp contractions to 20% of MVC have been widely used for estimating persistent inward currents (details below) [16,29,30]. To quantify force steadiness during 25% of MVC, the coefficient of variation of the force (CoV) was calculated = (SD/Mean)*100. To assess force tracking accuracy, all data were exported to Spike2 (version 9.11, CED Ltd., Cambridge, UK), where a virtual channel was created by subtracting the performed path from the requested (target) path and rectifying it. Area under the curve reflects the level of deviation from the target line reflecting muscle force tracking accuracy, with higher values indicating greater deviation.

### High-density surface electromyography

A semi-disposable high-density surface electromyography (HDsEMG) array (64 electrodes, 13×5, 8mm, I.E.D., GR08MM1305, OT Bioelettronica, Inc., Turin, Italy) was placed with the orientation of the muscle fibres (proximal to distal) over the muscle belly of right vastus lateralis after skin preparation involving shaving, light abrasion and cleansing with 70% ethanol. Electrodes were secured to the skin using flexible tape. The adhesive grids were attached to the surface of the muscle by disposable bi-adhesive foam layers (SpesMedica, Bettipaglia, Italy). The skin electrode contact was facilitated by filling the cavities of the adhesive layers with conductive paste (AC Cream, SpesMedica). A strap ground electrode (WS2, OTBioelettronica, Turin, Italy) dampened with water was positioned around the ankle of the right leg to optimise the signal quality. HDsEMG signals were acquired in a monopolar configuration, amplified (x256) with filtering set at 10-500 Hz and digitally converted at 2000 Hz by a 16-bit wireless amplifier (Sessantaquattro, OTBioelettronica, Turin, Italy) and transferred to a PC for further offline analysis. HDsEMG signals were recorded in OTBioLab software (OT Bioelecttronica, Turin, Italy).

The recorded HDsEMG signals were converted into a MatLab file comprising a single contraction. Each participant performed four submaximal contractions and ramp contractions; however, a single trial for each contraction protocol was selected for analysis based on the on the smoothness of the force profile. Monopolar HDsEMG signals were band-pass filtered at 20-500Hz and then decomposed offline into MU pulse trains by a convolutive kernel compensation algorithm [31,32]. After the decomposition, a trained investigator manually inspected the MU spike trains and edited the discharge patterns of each MU detected. Only MUs with a pulse-to-noise ratio equal or greater than 30 dB were kept for further analysis.

Mean discharge rate during the submaximal contraction at 25% of MVC was obtained and discharge rate variability was reported as the coefficient of variation (CoV) of the interspike interval (ISI) displayed as a percentage. Peak discharge rates during a ramp contraction were measured as the highest value of the smoothed MU discharge rate with a fifth order polynomial.

### Estimation of persistent inward currents (PICs)

The instantaneous firing rates of both MUs were calculated as the inverse of the interspike intervals of each MU spike train and smoothed by fitting a fifth order polynomial function. An estimate of the PIC magnitude was derived using the unit-wise MU analysis technique [17]. The lower threshold unit to be recruited in the ramp contraction is commonly referred to as the “control” unit; the “test” unit is a unit of higher threshold. A sufficient number of MUs were isolated for 10 young and 10 older participants. Delta F, the contribution of persistent inward currents to self-sustained firing, was calculated as the difference in “control” unit firing rate between the onset and offset of a “test” unit [16]. This technique relies on several assumptions: 1) test and control units share the common synaptic drive; 2) PIC is activated before or at the recruitment; 3) firing rate of the control MU closely reflects the depolarizing input to the parent motoneuron; 4) test and control unit process the synaptic inputs in a similar way. The MU pairs for the delta F calculation were selected based on the following criteria: 1) as a measure of common synaptic modulation, rate-to-rate correlation r ≥ 0.7; 2) to avoid the high variability in delta F calculation due to the initial acceleration phase of MU firings, test units were recruited at least 1s after the control units; 3) to account for the possibility of control unit saturation leading to an underestimation of delta F, pairs in which rate modulation of the control unit fell within 0.5pps were removed from analysis [16,30,33,34]. An average delta F was calculated when individual test units were paired with multiple control units (Figure 1).

**Figure 1.**
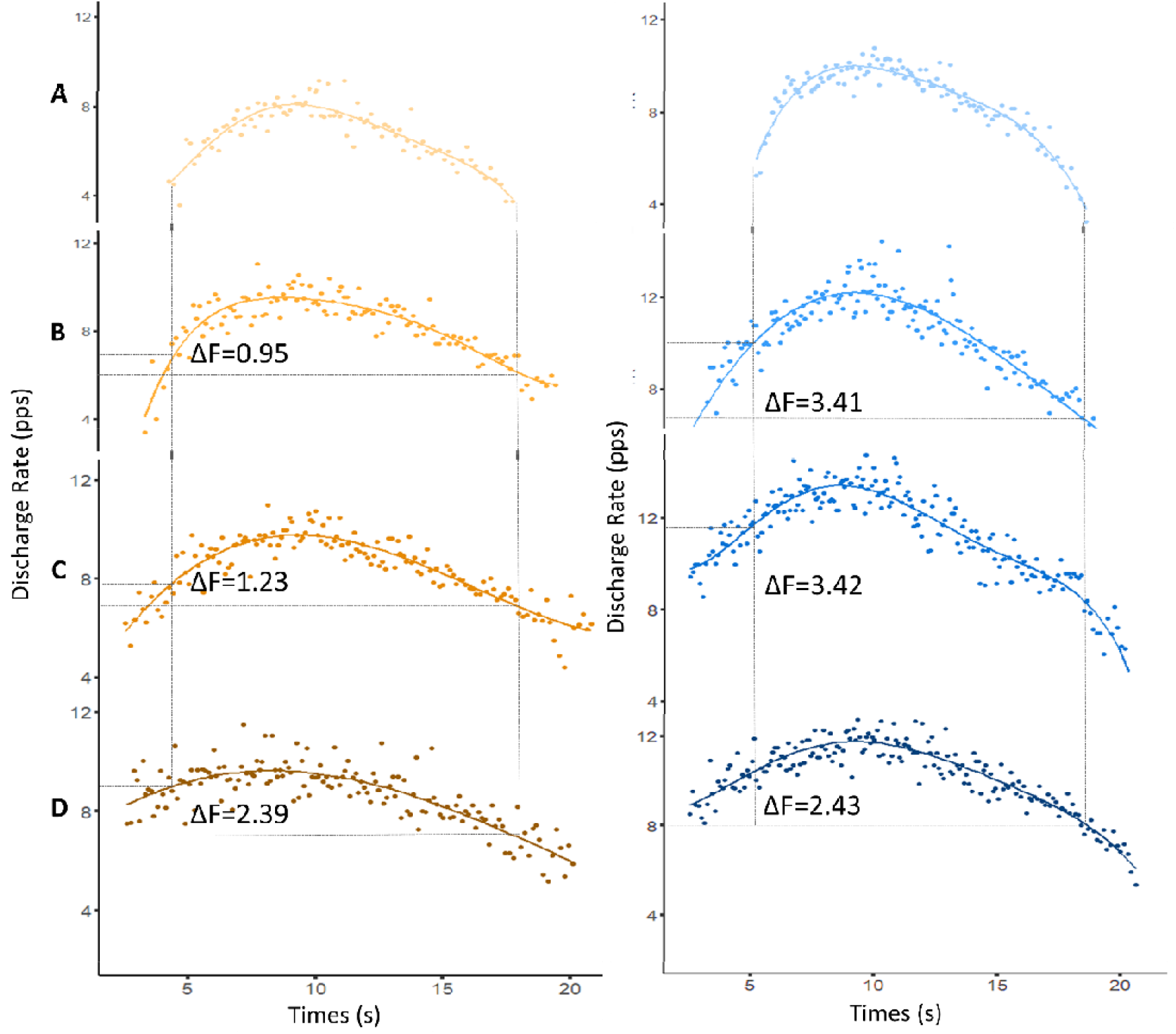
Example data showing delta frequency (ΔF) calculation in vastus lateralis for paired-motor unit technique. Paired motor unit data from a young male is displayed in the left panels (mean ΔF=3.09) and from an older male in the right panels (mean ΔF=1.52). Test units are displayed in panel A and corresponding control units are in B, C and D.

### Coherence Estimates

To estimate the level of common synaptic input, intramuscular coherence was calculated separately during sustained and ramp contractions (Figure 2), and represents a frequency domain correlation between cumulative spike trains (CSTs) of the identified motor units. The magnitude-squared coherence was calculated using the Welch’s averaged periodogram with 50% overlapping Hann windows of 1s at the different frequency bands: delta (0–5 Hz), alpha (5–12 Hz), beta (15–30 Hz) and piper (40–50 Hz) [35,36]. This analysis was repeated for 60 randomly chosen combinations of two equally sized CSTs calculated as the sum of firing times from three MUs randomly selected from the same set of MUs and then averaged. The Fisher Z-transform was applied to the coherence estimates to obtain a normally distributed variable for comparisons.

**Figure 2.**
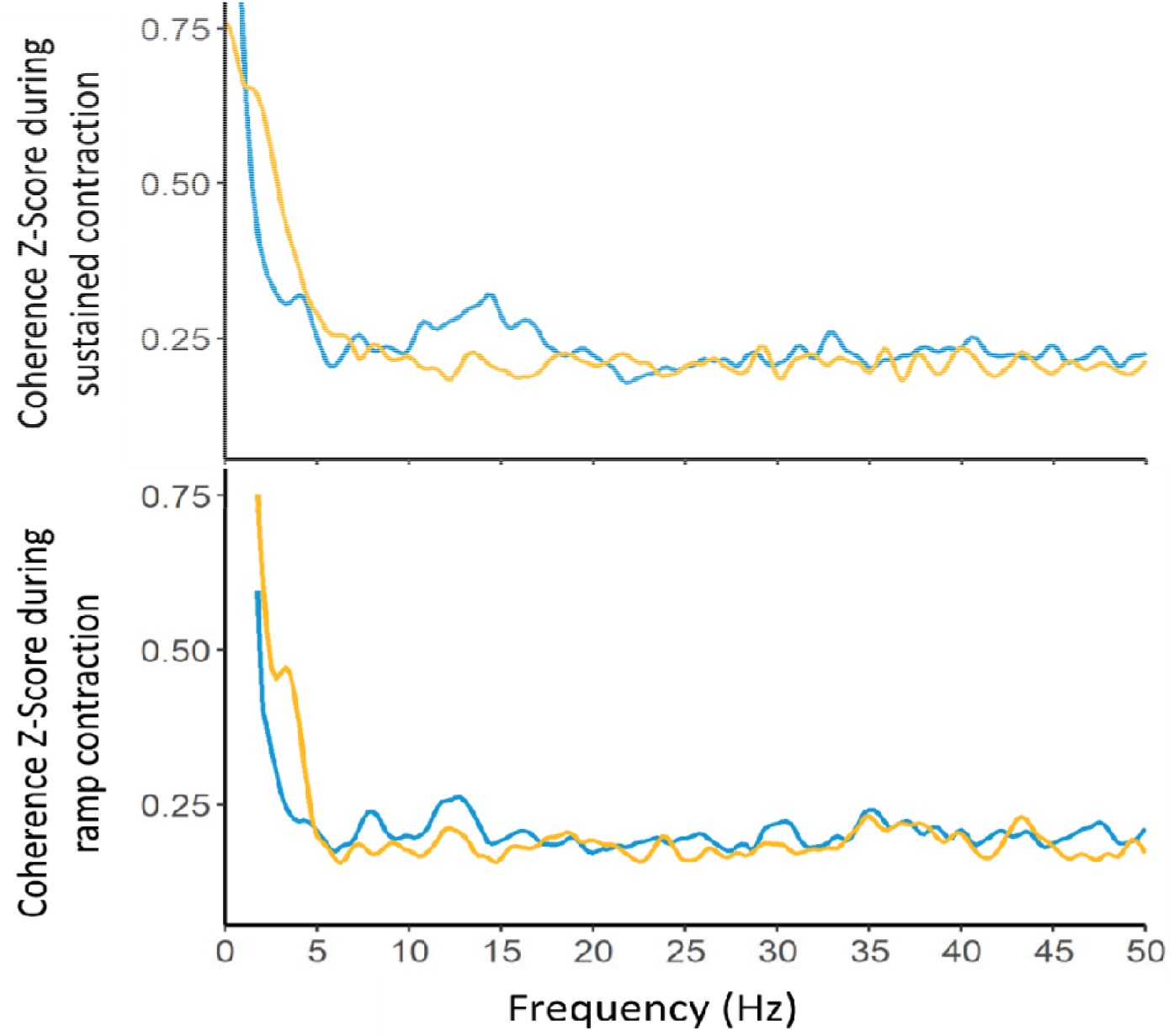
The group mean intramuscular coherence Z-Scores during submaximal contraction at 25% of maximal isometric voluntary contraction (MVC) and ramp contraction at 20% of MVC in young (blue) and older (yellow) participants. The magnitude-squared coherence was calculated from all the combinations (a maximum of 60 random permutations) of two equally sized cumulative spike trains, each composed of three motor units.

### Statistical analysis

Data management and analysis were performed using RStudio (Version 2022.07.1). Descriptive data were generated for all variables. *An independent t-test* was conducted to test whether any differences existed between young and older males in muscle cross-sectional area, torque, CoV-force, force tracking accuracy and coherence estimates. In order to preserve variability within and across participants simultaneously to the greatest extent, *multilevel linear regression models* were generated to compare mean discharge rate, discharge rate variability, peak discharge rate, delta F, recruitment threshold and derecruitment threshold between groups using *lme4 package* (Version 1.1–27.1) [37]. In the multilevel models, single MU was regarded as the first level; and individual participant with clustered MUs was considered as the second level. For data visualization, individual participant means are displayed in box-and-jitter plots. A *p* value <0.05 was considered statistically significant.

## Results

Young males had a larger muscle cross-sectional area (Y v O: 30.59 ± 5.87 v 23.34 ± 7.03 cm^2^, p=0.005; Figure 3A) and greater muscle torque (241.2 ± 62.69 v 143.3 ± 41.50 Nm, p<0.0001; Figure 3B) than older males. There was no difference in force steadiness (CoV-Force) (2.18 ± 0.73 v 2.27 ± 0.65, p=0.735; Figure 3C) during sustained contractions. However, there was a significant difference between groups in force tracking accuracy of the ramp contraction, with the younger group performing better than the old (9.41 ± 3.25 v 19.47 ± 8.36 Ns, p<0.001; Figure 3D).

**Figure 3.**
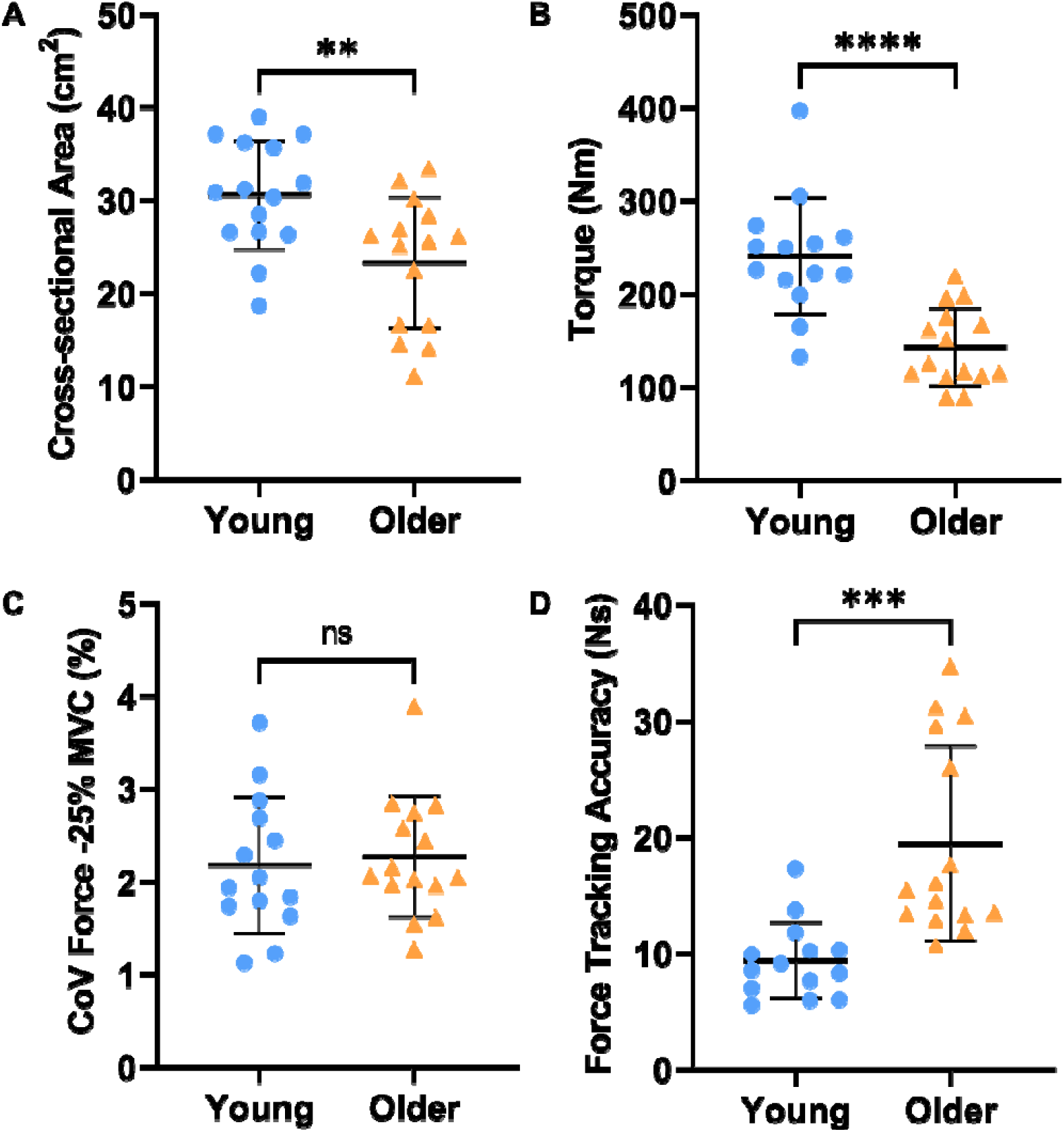
Cross-sectional area (A), torque (B), coefficient of variance of force at 25% maximal voluntary contraction (C) and force tracking accuracy (area under curve) during a single ramp contraction at 20% of maximal voluntary contraction (D) between young (blue) and older (yellow) males. Lines indicate group means and SD. ***p<0.001, **p<0.010, *p<0.05.

During the submaximal contraction at 25% of MVC, a total of 246 MUs were identified from young males and 282 MUs from older males. There was no statistical difference in mean discharge rates (8.50 ± 2.08 v 7.98 ± 1.64, p=0.137, Figure 4A) or discharge rate variability (11.34 ± 6.01 v 10.22 ± 5.15, p=0.072, Figure 4B) between groups.

**Figure 4.**
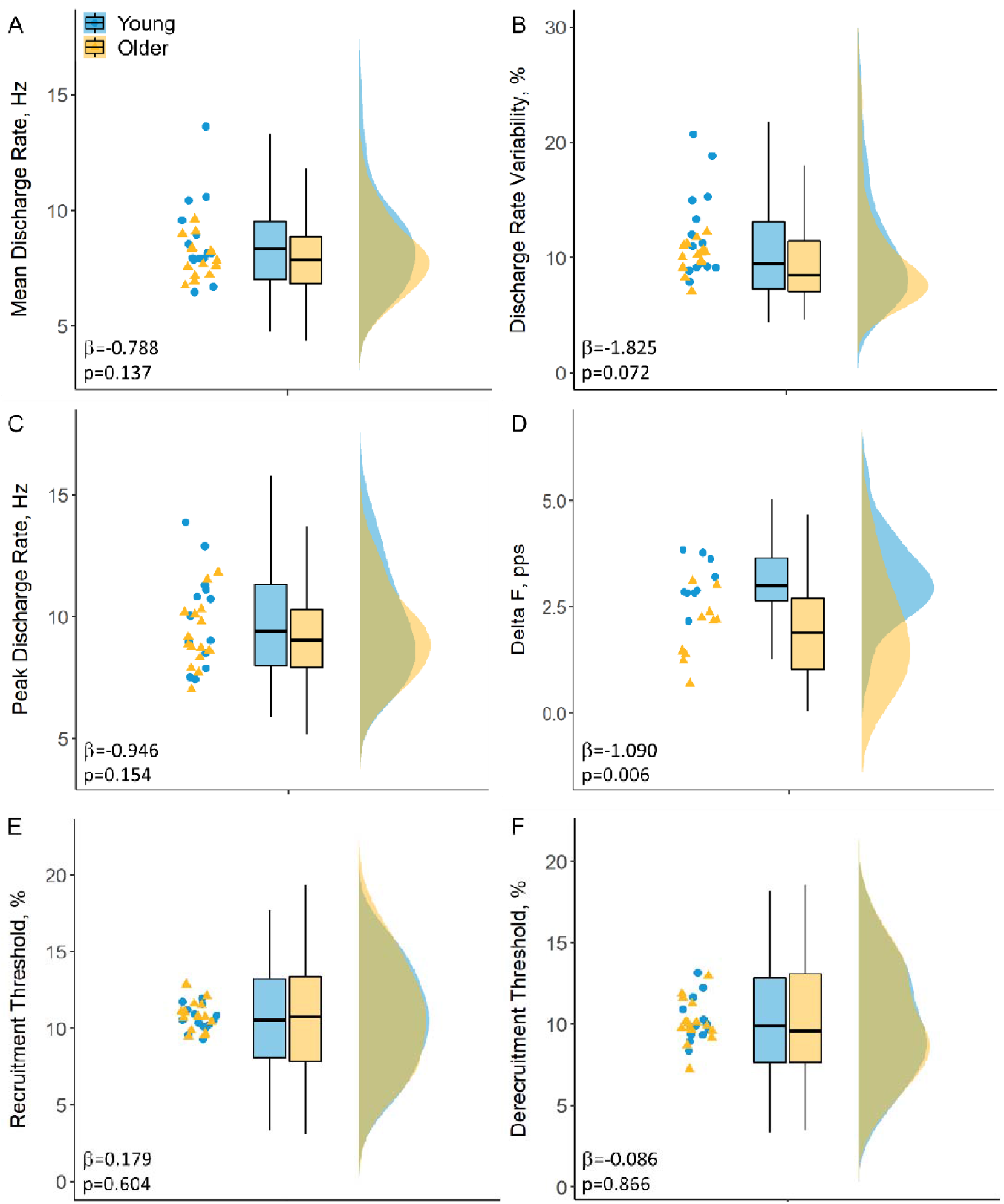
Jitter plots of the individual participant mean values and box and raincloud plots of mean discharge rate (A), discharge rate variability (B), peak discharge rate (C), Delta F (D) and recruitment/decruitment threshold (E&F) in young (blue) and older (yellow) males.

During the ramp contractions peaking at 20% of MVC, a total of 256 MUs were identified from young males and 262 MUs from older males. There was no statistical difference in peak discharge rates (9.63 ± 2.20 v 9.35 ± 1.94 pps, p=0.154; Figure 4C). For Delta F calculations, there was an average of 5 (SD: 3.68) pairs of test and control units per person in the young and 6 (4.95) pairs in older group, with the mean number of 4 (1.79) test units per person in young and 3 (1.97) in old. Younger males had significantly greater delta F values when compared to older participants (3.11 ± 0.77 v 1.86 ±0.65 pps, p=0.006; Figure 4D). There was no significant difference in recruitment (10.60 ± 3.37 v 10.75 ± 3.55 %, p=0.604; Figure 4E) or derecruitment thresholds (10.14 ± 3.40 v 10.18 ± 3.39 %, p=0.866; Figure 4F) of the identified MUs between young and older groups.

In the sustained contraction, there was no significant age-related difference in coherence in Delta (0.49 ± 0.14 vs 0.54 ± 0.11, p=0.315), Alpha (0.24 ± 0.05 vs 0.23 ± 0.03, p=0.462) or Beta (0.22 ± 0.03 vs 0.21 ± 0.02, p=0.125) bands, however the young had higher coherence in the Piper band (0.23 ± 0.03 vs 0.21 ± 0.02, p=0.026) (Figure 5A-D). In the ramped contraction, there were no age-related differences in COH Delta (0.87 ± 0.08 vs 0.92 ± 0.12, p=0.182), Beta (0.19 ± 0.02 vs 0.18 ± 0.02, p=0.110) or Piper (0.20 ± 0.04 vs 0.18 ± 0.02, p=0.156) bands however, young participants had greater coherence estimates in Alpha band (0.21 ± 0.02 vs 0.18 ± 0.01, p<0.001) (Figure 5E-H).

**Figure 5.**
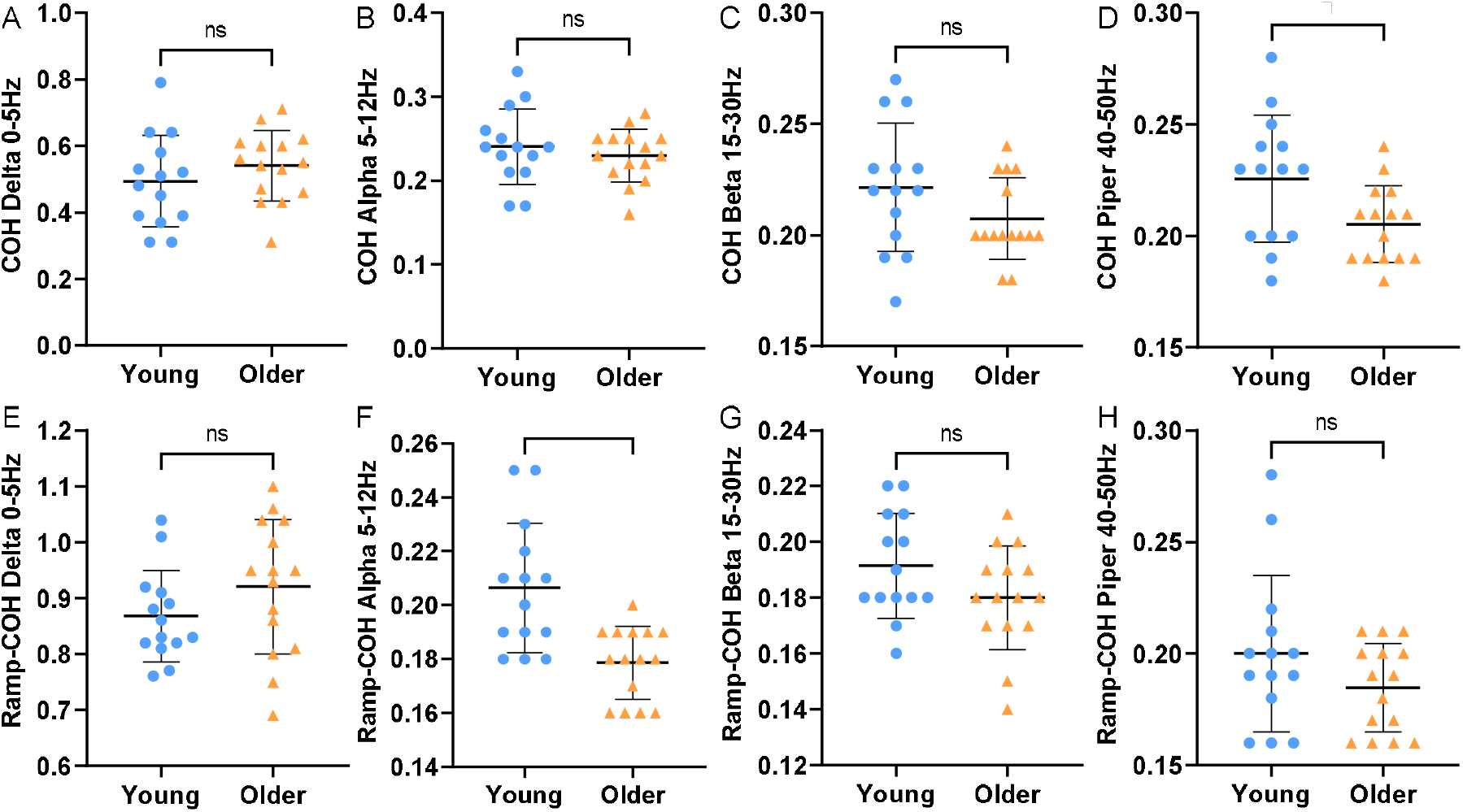
Individual participant means of intramuscular coherence Z-score at different frequency bands during a submaximal contraction at 25% maximal voluntary contraction (MVC) and a ramp contraction at 20% MVC in young (blue) and older (yellow) males.

## Discussion

Here we provide the first evidence for reduced estimates of persistent inward currents combined with reduced common synaptic inputs alongside poorer force control ability in the aged human VL, a muscle with established susceptibility for age-related functional losses. Though no significant age-related difference in MU discharge rates was observed, our findings highlight important aspects of impaired neural input to older muscle that likely contribute to functional decrements.

A smaller muscle size and a lower muscle strength were observed in older participants, which is consistent with the wider ageing literature [38,39]. However, no age-related difference in MU discharge rates was detected between groups in both submaximal sustained contraction or ramp contraction, which is not aligned with our previous findings in this muscle using needle EMG recordings [39]. This may reflect differences in the population of MUs sampled, with needle EMG sampling from a greater depth of muscle than surface-based HD EMG used here [40].

Age-related differences in force control ability were only observed during the ramp contraction, highlighting the role of task complexity in age-related impairments [41]. Numerous factors associated with advancing age may also explain this poorer force control in a more complex contraction, including MU remodelling, reduced excitability of the MUs, decreased muscle spindle sensitivity, less common synaptic inputs, and/or impaired cutaneous afferents [42,43]. Estimates of MU number in the VL are reduced in older age [39,44], and although direct human evidence is lacking, animal models show large MUs are preferentially lost with ageing and denervated fibres are reinnervated by early recruited MUs [45], resulting in expansion of those earlier recruited MUs [46] such as those sampled in the current study (< 20% MVC). These MUs with a larger number of fibres, when activated, would have a greater influence on force during their progressive re/derecruitment processes, and may partly explain the age-related differences of force control observed here.

The progressive increase and decrease in force during the targeted isometric ramped contraction used here is partly reliant on afferent neuromodulatory feedback from the site of force application [47]. Several studies have reported a decrease of grey matter and white matter throughout the motor cortex with increasing age [48–50] and a strong correlation between cortical atrophy and fine control capacity [51]. A significant decrease in inhibition has also been observed in aged muscles, as well as a greater level of cortical and subcortical area activity during a targeted motor control task [52–54], all of which may exert greater detrimental effects during a complex task than simple (e.g. ramped vs sustained). Additionally, both animal and human studies have revealed a decreased proprioception in aged muscles [55] with the evidence of declined muscle spindle numbers and degeneration of the sensory nerve terminals, leading to the morphological adaptations at a peripheral level and modulation of mechanoreceptor gain at a central level [56,57]. Decreased inhibitory spinal circuits with ageing has also been shown to have influence on Ia afferents from muscle spindles to spinal motoneurons [58].

Although not directly assessed, the points made thus far are supported by the intramuscular coherence findings during the ramp contraction. Coherence in alpha band (5-12 Hz) is associated with Ia afferent feedback [59] and when exploring the age-related differences, a significant lower coherence estimate in alpha band was observed in old only during the ramp contraction, corresponding to the poorer force tracking ability. This result supports evidence from previous studies, showing a greater loss of Ia afferent feedback [60] and an impaired ability to modulate Ia presynaptic inhibition [61] in older participants. Moreover, it has also been suggested that age-related deterioration of cutaneous afferent inputs may also contribute to reduced force control ability [62]. Therefore, a decreased ability to regulate the progressive increase and decrease in force may also be a result of the combination of reduced Ia afferent feedback from muscle spindles and reduced cutaneous afferents.

The effect of ageing on MU recruitment thresholds is inconclusive, or more accurately, it appears to be muscle specific. In a single study of the biceps and triceps, MUs were recruited earlier in older triceps, but not the biceps [17]. This study of VL found no age-related difference in mean MU recruitment or derecruitment threshold, which was approximately 10% of MVC in both young and older participants during contractions up to 20% of MVC; indicating age was probably not the main predictor of MU recruitment/derecruitment threshold of VL.

Compared to younger participants, we report a significantly reduced delta F in the VL of older participants during a ramp contraction, supportive of studies of other muscles in the upper [17] and lower [18] limbs. The current findings in VL showed that the estimates of PICs were lower in old, which is likely attributable to the reduced activity of the monoaminergic system. Serotonin (5-HT) and norepinephrine (NE) appears primarily to be involved in the modulation of motor control and sensory input, partly influencing the magnitude of PICs [63]. Ageing is associated with decreased levels of 5-HT receptors and transporters in the brain [64,65] as well as impaired NE synthesis and secretion [66], leading to a reduced monoaminergic neurotransmission. Additionally, the magnitude of PICs may also be influenced by age-related alterations in Na^+^ and L-type Ca^2+^ ion channels, widely distributed on the dendrites of motoneurons. The dysregulation of Ca^2+^ has been reported to have an essential role in the development of ageing and the neurodegeneration process via C. elegans models [67], which may also be apparent in human spinal motoneurons. A high sensitivity of PICs to inhibitory input to motoneurons has been reported [12] and as discussed above, inhibitory inputs significantly decrease with advancing age at the brain and spinal cord levels.

### Strengths/Limitations

To our knowledge, this is the first study concurrently investigating the upper motor neuron command (common synaptic inputs) and lower motor neuron excitability (persistent inward currents) in young and older VL. Age-related impairments are evident in each. However, there are several limitations. Firstly, the results reported here were from males only which may not directly translate to females, as decreases in estrogen levels may also contribute to the alterations in circulating serotonin and its receptor densities in postmenopausal women [68]. Secondly, the MU data we report are relevant to those recruited at lower contraction levels (< 25%) and reveal little of those later recruited MUs. In addition, we used 12s and 20s signal length for calculating coherence estimates during submaximal sustained contraction and ramp contraction respectively, possibly resulting in an underestimated effect of ageing on common synaptic inputs.

## Conclusion

The reduced muscle strength and control ability observed in older males is partially attributable to reduced estimates of persistent inward currents and common synaptic input, which occur independently of reductions in MU discharge rates. These findings have important implications in the field of healthy human ageing and should be considered when applying interventions to mitigate age-related functional decrements.

## Data Availability Statement

The datasets generated and analysed during the current study are available from the corresponding author upon reasonable request.

